# Three-step docking by WIPI2, ATG16L1 and ATG3 delivers LC3 to the phagophore

**DOI:** 10.1101/2023.07.17.549391

**Authors:** Shanlin Rao, Lisa M. Strong, Xuefeng Ren, Marvin Skulsuppaisarn, Michael Lazarou, James H. Hurley, Gerhard Hummer

## Abstract

The covalent attachment of ubiquitin-like LC3 proteins prepares the autophagic membrane for cargo recruitment. We resolve key steps in LC3 lipidation by combining molecular dynamics simulations and experiments *in vitro* and *in cellulo*. We show how the E3-like ligase ATG12– ATG5-ATG16L1 in complex with the E2-like conjugase ATG3 docks LC3 onto the membrane in three steps by (1) the PI(3)P effector protein WIPI2, (2) helix α2 of ATG16L1, and (3) a membrane-interacting surface of ATG3. Phosphatidylethanolamine (PE) lipids concentrate in a region around the thioester bond between ATG3 and LC3, highlighting residues with a possible role in the catalytic transfer of LC3 to PE, including two conserved histidines. In a near-complete pathway from the initial membrane recruitment to the LC3 lipidation reaction, the three-step targeting of the ATG12–ATG5-ATG16L1 machinery establishes a high level of regulatory control.

## Introduction

Eukaryotic cells use autophagy for the wholesale degradation of bulk cytosol and bulky substrates, including intracellular pathogens, protein aggregates, and mitochondria^1^. Autophagy of the latter is referred to as mitophagy^2, 3^. Defects in mitophagy downstream of the E3 ubiquitin (Ub) ligase Parkin and the Ub kinase PINK1 are implicated in familial Parkinson’s disease^4^. Autophagy is critical for cell homeostasis across a vast range of physiological conditions, and its defects contribute to essentially all the major late-onset neurodegenerative diseases, cancer, and other diseases^5^. The covalent conjugation of the ubiquitin-like ATG8 proteins to the membrane lipid phosphatidylethanolamine (PE) is a hallmark of autophagy^6^. Atg8 is the sole and founding member of this family in yeast, and it has six orthologs in humans, LC3A-C, GABARAP, and GABARAP L1-2^7^. ATG8 family proteins bind to short motifs known in humans as LC3-interacting regions (LIRs). LIR motifs are found throughout the machinery of autophagy, where their interactions facilitate cargo sequestration in selective autophagy^8^, autophagosome-lysosome fusion and autophagic membrane breakdown^9, 10^, and indeed, have some role in most steps in autophagy.

ATG8s are conjugated to membrane PE through a pathway that has both analogies and differences with protein ubiquitylation. Ub and Ub-like proteins are conjugated to proteins, usually via the ε-amino group of Lys residues, by the sequential action of E1, E2, and E3 enzymes^11, 12^. The chemistry and structural biology of protein ubiquitylation, and the related Neddylation, SUMOylation, and similar pathways, have been elucidated in great detail^11, 12^. ATG8 conjugation begins with the action of the E1-like ATG7 and the E2-like ATG3 enzymes^6^. These enzymes have the same overall fold and active site cysteine residue as their cognate Ub E1 and E2 enzymes^13^, as well as unique modifications that facilitate their mutual interaction^14^ and the interaction of ATG3 with membranes^15^. Purified ATG3 can carry out ATG8ylation on highly curved liposomes *in vitro*^15^ in the absence of its cognate E3, but *in vivo*^16^ and even in a giant unilamellar vesicle (GUV) reconstitution system^17, 18^, the downstream E3 complex components are essential.

The autophagic counterpart of the Ub E3 is the ATG12–ATG5-ATG16L1 complex^19^, which is structurally and evolutionarily unrelated to any of its functional equivalents in ubiquitylation. The ATG12-ATG5 unit is itself covalently bonded through an ATG10-dependent reaction^20^. ATG12-ATG5 binding allosterically activates ATG3 by increasing the exposure and reactivity of its Cys264-linked ATG8 thioester for transfer to PE^14, 21^. The ATG16L1 portion of the complex is responsible for delivery and positioning on the membrane^19^. ATG16L1 is itself delivered to membranes by the β-propeller protein WIPI2^18, 22^. WIPI2 (and other WIPIs) are recruited to membranes early in autophagy induction by the lipid phosphatidylinositol 3-phosphate (PI3P)^23^, which is generated by the class III PI 3-kinase complex I (PI3KC3-C1) early in autophagy initiation^24^.

The problem of how the chemistry and structural biology of a protein ubiquitylation-like system is adapted to act on a membrane substrate has been one of the major open questions in the mechanistic biochemistry of autophagy. A number of pieces of the puzzle have come together in recent years. The structural basis of the assembly of a fragment of ATG3 with the ATG12–ATG5-ATG16L1 unit was worked out for the human proteins^21^. ATG16L1 contains an amphipathic helix ɑ2, adjacent to its ATG5 binding site, that is strongly sensitive to membrane curvature^25^ and essential for promotion of LC3 lipidation in liposomes and in cells^26^. It is puzzling that ATG16L1 ɑ2 is so important for catalysis, given that WIPI2 is capable of recruiting ATG16L1 to membranes through its WIPI2-interacting region (W2IR)^18, 22^. The structural basis for human ATG16L1 recruitment by WIPI2 has also been worked out^18, 27^. The ATG12-ATG5 and WIPI2-binding regions of ATG16L1 are separated by a coiled coil with > 100 amino acids. The resulting extended shape and its flexibility challenge experimental structure determination of the full membrane bound WIPI2-ATG12–ATG5-ATG16L1-ATG3 system. Here, we approached the problem beginning with large-scale all-atom simulations of the WIPI2-ATG12–ATG5-ATG16L1-ATG3 on lipid membrane. Predictions from the simulations were verified experimentally *in vitro* and *in cellulo*. In this way, we connect structural and biochemical information into a near-complete view of the lipidation pathway.

## Results

### Docking step 1: WIPI2 recruits ATG12–ATG5-ATG16L1 loaded with ATG3-LC3 to phagophore

As a key first step in targeting the lipidation machinery to the phagophore membrane, we concentrated on the WIPI2-mediated membrane interaction of ATG12–ATG5-ATG16L1. The central homodimer-forming coiled-coil domain (residues 78-230) of the human ATG16L1 protein is predicted^28^ to form a continuous stretch of α-helical coiled coils spanning the major part of the domain (∼115 amino acids from the N-terminal side), allowing reconstruction of the dimeric ATG16L1 structure by fitting geometric parameters^29^ based upon Crick’s equations^30^ (SI Fig. 1A). The resulting coiled-coil structure is in excellent agreement with crystal structures^31, 32^ of the mouse orthologue in which an overlapping region of the coiled-coil domain has been resolved (SI Fig. 1A), providing validation for our ATG16L1 model. Using AlphaFold^33, 34^, we also obtained a structural model of the E2-like ATG3 conjugase loaded with LC3 (SI Fig. 1B). The predicted ATG3-LC3 complex adopts a conformation compatible both with binding to the E1-like ATG7 homodimer in the preceding step and with formation of a thioester bond between the catalytic Cys264 side chain of ATG3 and the C-terminal Gly120 of LC3 to yield the E2-substrate conjugate (SI Fig. 1B). The core of the human ATG3 structure is architecturally similar to the yeast and *Arabidopsis* Atg3 proteins, as has been previously reported^35^, with an intrinsically disordered region^36^ forming an ∼100-residue loop that contains the ATG12-binding sequence^21^ (Fig. 1A) as well as a region predicted to participate in β-sheet formation in the presence of LC3. Combined with crystallographic structures^21, 37^ of ATG12-ATG5 in quaternary complex with a bound fragment of ATG3 and the N-terminal ATG5-binding domain of ATG16L1, we present an atomistic model of the full LC3 lipidation machinery consisting of the E3-like ATG12–ATG5-ATG16L1 complex bound to the E2-substrate conjugate, ATG3-LC3 (Fig. 1A).

**Figure 1.**
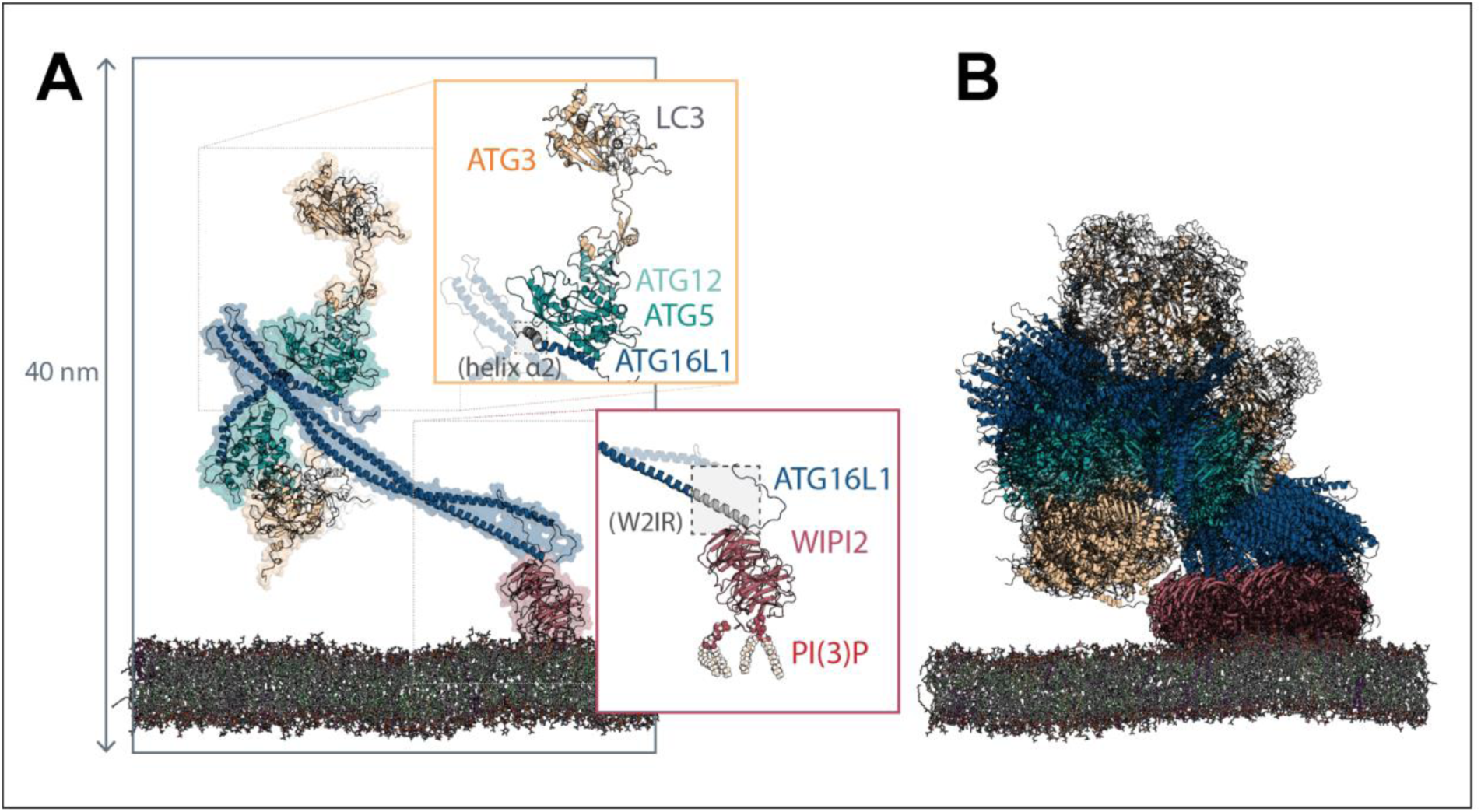
Structure and dynamics of the membrane-recruited ATG12–ATG5-ATG16L1-WIPI2 complex loaded with ATG3-LC3. (**A**) Ribbon and (semi-transparent) surface representation of the full LC3 lipidation machinery bound to a membrane mimicking the phagophore lipid composition, upon equilibration and atomistic molecular dynamics simulation. The ATG16L1 N-terminal helix α2 and the WIPI2-interacting region (W2IR) are highlighted in gray. (**B**) Dynamics of the assembly, illustrated with a superimposition of conformations sampled at 50 ns intervals during the final 500 ns of one 1 µs simulation trajectory. Flexibility of inter domain loops allows the ATG3-LC3 conjugate (yellow/white) to explore the region of space above the membrane to which the ATG12–ATG5-ATG16L1 is anchored.

To determine the configuration of the ATG12–ATG5-ATG16L1 complex recruited to phagophore membranes by the PI(3)P effector WIPI2, we first performed atomistic molecular dynamics (MD) simulations on a WIPI2-ATG16L1 co-crystal structure^18^ in which WIPI2 is bound to the WIPI2 interacting region of ATG16L1 (residues 207-230). Initially placed at a minimum distance of ∼2 nm above PI(3)P-containing membranes mimicking the ER lipid composition^38^, WIPI2 formed spontaneous membrane contacts in an expected orientation, with the two putative phosphoinositide binding sites^39–41^ in its β-propeller blades 5 and 6 interacting with PI(3)P (Fig. 1A) and the N-terminal side of the bound ATG16L1 segment oriented away from the membrane. By aligning our structural model of ATG12–ATG5-ATG16L1 described above to the membrane-associated WIPI2-ATG16L1 configuration established during simulations, a first model of the membrane recruited LC3 lipidation machinery was thus obtained.

Upon atomistic MD simulations of ATG12–ATG5-ATG16L1 complexed with the ATG3-LC3 conjugate and anchored via WIPI2 to membranes approximating the phagophore lipid composition, all components maintained structural integrity across all five 1 µs simulation replicates (SI Fig. 2). Flexing and tilting motions of the dimeric coiled-coil structure of ATG16L1 were accompanied by considerable flexibility exhibited in the inter-domain loop regions of ATG16L1 and ATG3 (Fig. 1B), allowing the ATG3-LC3 conjugate to explore favorable binding configurations near the membrane (SI Movie 1). However, membrane binding was observed only in the case where ATG3-LC3 was already at the membrane surface upon initiation of the simulation. This finding indicates that the upward tilt of the WIPI2-attached coiled coil tends to keep the ATG3-LC3 conjugate above the membrane, even if direct interactions of ATG3-LC3 are possible in principle.

**Figure 2.**
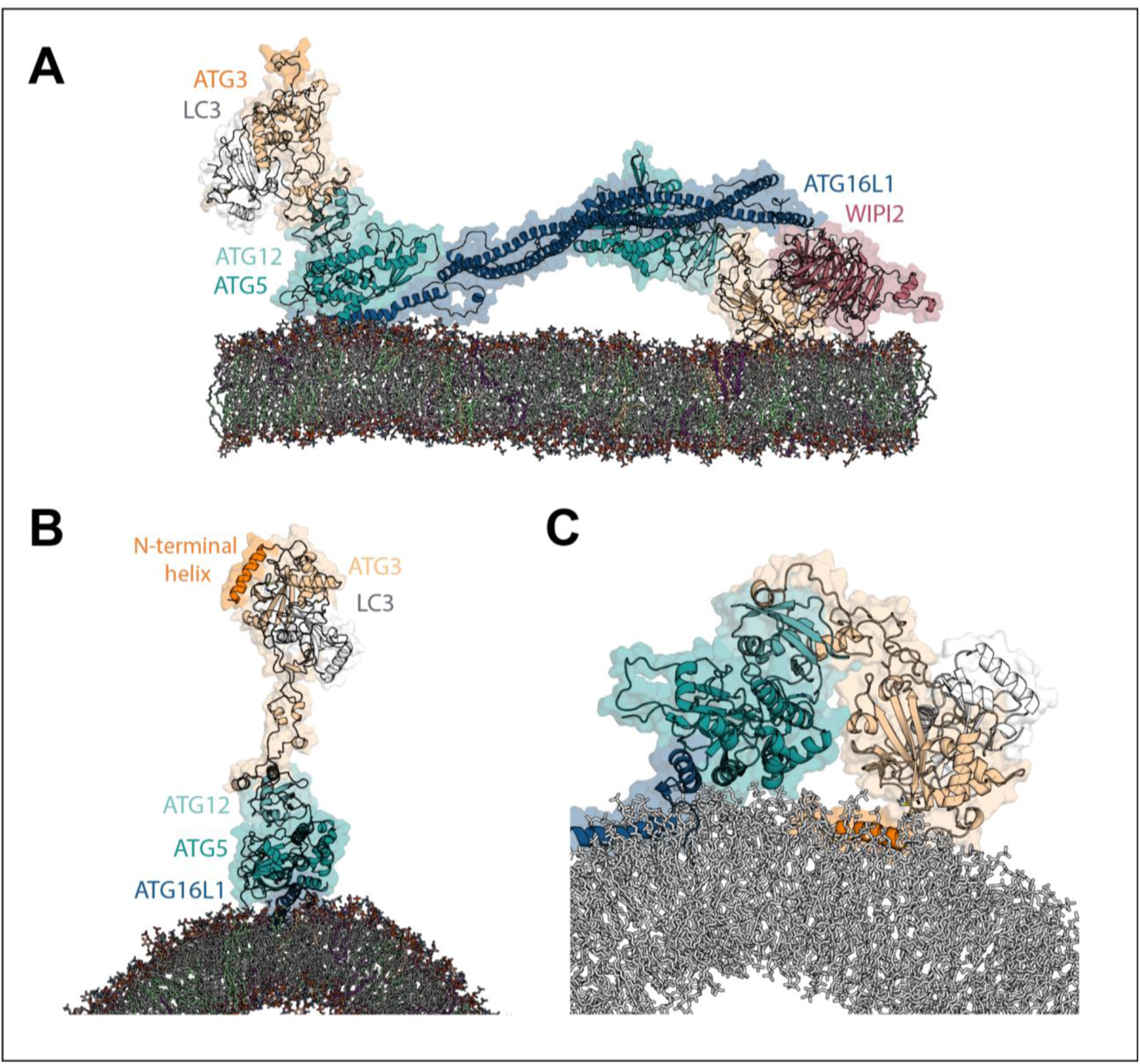
Membrane-bound ATG16L1 helix α2 brings ATG3-LC3 into membrane contact. (**A**) Snapshot of the WIPI2-recruited ATG12–ATG5-ATG16L1 complex with the N-terminus of ATG16L1 also engaged in membrane interaction in an atomistic molecular dynamics simulation. The ATG16L1 helix α2 had been steered gently to the membrane surface at a speed of 0.5 nm ns^-1^, with the complex subsequently relaxed in extended (1 μs) simulations. (**B**) Initial configuration of simulations representing a further stage of ATG3-LC3 delivery by ATG12–ATG5-ATG16L1, following membrane recruitment via WIPI2. The hydrophobic face of helix α2 is embedded into the membrane at this stage. (**C**) Snapshot of ATG3-LC3 delivered to the membrane whilst bound to ATG12–ATG5-ATG16L1, with the ATG3 N-terminal helix (orange) spontaneously inserted between membrane lipids. Taken at *t* = 450 ns from a 1 μs simulation replicate.

To reconcile the prevailing model of membrane recruitment of ATG12–ATG5-ATG16L1 by WIPI2 with the requirement for the membrane-interacting ATG16L1 N-terminal helix α2^26^, we hypothesized that upon initial recruitment through WIPI2, direct membrane binding by helix α2 constitutes a crucial second step in delivering ATG3-LC3 nearer to the target membrane. We focus here on the *cis* configuration in which the entire LC3 lipidation machinery becomes associated with the same patch of membrane^42^. However, our molecular model does not rule out the alternative possibility whereby ATG12–ATG5-ATG16L1 anchored at omegasomal membranes would bridge an intermembrane distance to facilitate LC3 conjugation to the nascent phagophore in *trans*^22^.

### Docking step 2: Helix α2 of ATG16L1 pulls ATG3-LC3 to membrane

As a possible second step in membrane targeting, we focused on helix α2 of the ATG16L1 N-terminal domain, which has been shown to bind membranes^26^ with a preference for positive membrane curvature^25^. For an ATG12–ATG5-ATG16L1 complex attached to the membrane via WIPI2, this mode of membrane interaction is made possible by the flexibility of the ∼30-residue inter-domain loop between the N-terminus of ATG16L1 and its coiled coil. We demonstrated this ability of ATG16L1 to engage with the membrane simultaneously, at one end, through recruitment by WIPI2 and, at the other end, via helix α2 by gently pulling the ATG16L1 helix α2 toward the membrane in steered MD simulations and then relaxing the complex in extended MD simulations (Fig. 2A).

Building upon our previous MD simulations of ATG12–ATG5-ATG16L1 binding to curved membranes^25^, with ATG16L1 interacting either at the membrane surface or with an embedded hydrophobic face of the amphipathic helix α2, we obtained models of membrane-bound ATG12–ATG5-ATG16L1 loaded with ATG3-LC3 (Fig. 2B). In atomistic MD simulations, the flexible ATG3 loop then allowed ATG3-LC3 to reach the membrane spontaneously whilst maintaining its interactions with the α2-anchored ATG12–ATG5-ATG16L1 complex (SI Movie 2). In this configuration, we found the N-terminal helix of the ATG3 conjugase to embed into the membrane (Fig. 2C and SI Movie 2) without any biasing force. Once inserted, the ATG3 amphipathic helix remained embedded in the membrane through the course of the simulation, establishing stable membrane contacts. This finding is consistent with the previously reported role of the ATG3 N-terminus as an essential membrane-targeting element^43^. Geometrically, the two ATG3-LC3 conjugates flexibly connected to the ATG12–ATG5-ATG16L1 complex can simultaneously engage with the membrane for parallel lipidation reactions.

### Docking step 3: Catalytic domain of ATG3 forms stable membrane interaction interface upon membrane insertion of ATG3 N-terminal helix

For a decisive third and final targeting step, we explored how the catalytic domain established membrane contact. The ATG3 conjugase has been reported to show basal activity *in vitro* for catalyzing LC3 conjugation in the absence of ATG12–ATG5-ATG16L1^21^. Having observed stable membrane association of the ATG3-LC3 conjugate held near phagophore-mimetic membranes by ATG12–ATG5-ATG16L1, we sought to further collect lipid contact data on ATG3-LC3 by initiating a set of longer (2 µs) replicates of smaller simulation systems containing the isolated conjugate placed directly above membranes. In eight of the twenty trajectories thus obtained, spontaneous membrane insertion of the ATG3 N-terminal helix occurred within the first 1 µs. Comparison of ATG3-LC3 lipid contacts (post-insertion of the ATG3 helix) reveals a consistent membrane interaction interface in presence or absence of ATG12–ATG5-ATG16L1 (Fig. 3A). We found that the ATG3 protein dominated the interactions of the conjugate with PE-containing membranes.

**Figure 3.**
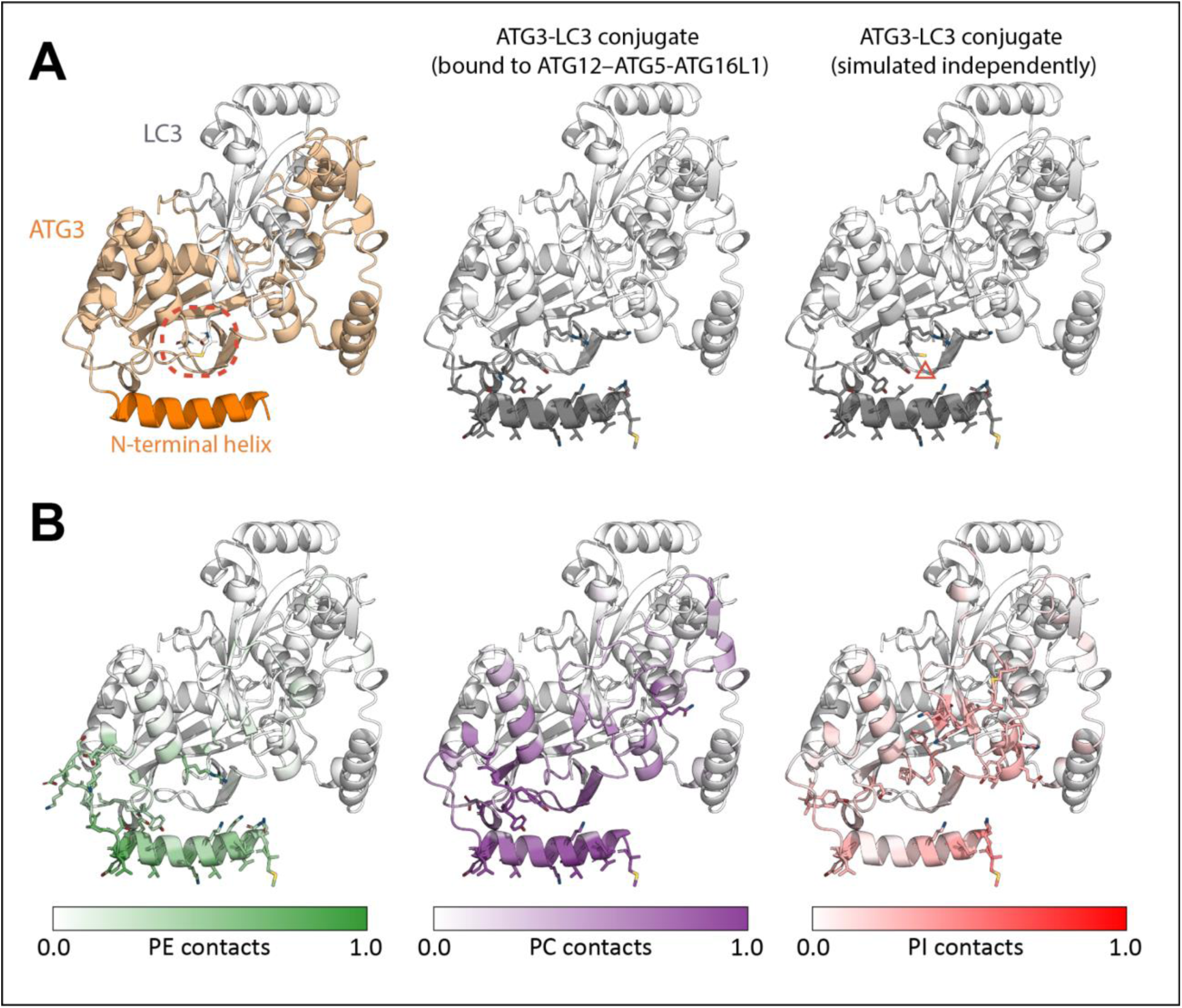
Membrane lipid contacts formed by the ATG3-LC3 conjugate. (**A**) Ribbon representation of the ATG3-LC3 conjugate structure prior to simulation, with the catalytic site indicated with a red dashed circle around the thioester bond (left). Coloring ATG3-LC3 residues by their mean frequency of membrane contacts (middle/right; white to gray at increasing contact frequency), upon spontaneous insertion of the ATG3 N-terminal helix in atomistic molecular dynamics simulations, reveals a consistent membrane interaction interface whilst bound to ATG12–ATG5-ATG16L1 (final 500 ns of a 1 μs trajectory) and in the absence thereof (final 1 μs of each of eight 2 μs replicates). Top-ranked residues are highlighted as sticks: the only qualitatively discernible difference between membrane interaction data from the two independent simulation systems is indicated with a red triangle. (**B**) Proportion of membrane contacts formed by ATG3-LC3 residues with different types of lipids present in the membrane, illustrated with phosphatidylethanolamine (PE), phosphatidylcholine (PC), and phosphatidylinositol (PI). Membrane interaction data are averaged across the final 1 μs of eight 2 μs replicates and normalised for each lipid type such that a value of 1.0 is assigned to the residue(s) showing highest specificity for that lipid.

### ATG3-LC3 presents active site towards the membrane in configuration conducive to lipidation reaction

With ATG3-LC3 at the membrane, we explored the structural foundation of the actual lipidation reaction. The folded core of ATG3 comprises a six-stranded β-sheet (strands β1-β6) surrounded by α-helices^44^. Among regions of ATG3-LC3 that formed frequent membrane interactions in our simulations were short sequences of residues within inter-secondary structure loops of the ATG3 core, namely (i) catalytic domain residues 208-211 and 242-243 of the β3/β4 and β4/β5 loops, respectively; (ii) residues 262-265 encompassing the thioester-forming Cys264 between β6 and the succeeding α-helix; and to a lesser extent (iii) residues 61-64 within the β1/β2 loop. The catalytic site, which contains Cys264 of ATG3 covalently bonded to the C-terminus of LC3, was situated centrally on the membrane interaction interface identified above and exposed towards membrane lipids (Fig. 3A). Furthermore, the ATG3-LC3 conjugate formed distinct interactions with different types of lipids present in the membrane, with PE localizing particularly near the catalytic center (Fig. 3B). Our data thus suggest a preferred orientation of ATG3-LC3 on the membrane that is compatible with catalyzing LC3 conjugation to the phagophore.

### Mutational analysis and functional validation

The MD simulations identified a surface of ATG3 that was consistently in contact with the membrane in the context of the larger ATG12–ATG5-ATG16L1-ATG3–LC3B-WIPI2 complex. Residues in this patch include Lys62, Lys64, Lys208, Tyr209, Tyr210, Thr244, His262, Cys264, Arg265, and His266 (Fig. 4A). The presence of Cys264 was expected, given this residue’s known role as the LC3 donor in the reaction^21, 45, 46^. We assayed LC3B conjugation activity in a small unilamellar vesicle (SUV) system similar to that originally used to demonstrate Atg8 conjugation activity of the Atg12–Atg5-Atg16-Atg3 complex^46^. Here, purified human proteins were used (1.0 μM), WIPI2 (0.5 μM) was included in the protein mixture, and 10% PI(3)P included in the SUVs^18^. Activity was monitored by the conversion of LC3B-I to LC3B-II. As expected, essentially complete conversion was seen for wild-type, while the mutation C264A of the catalytic cysteine as a negative controlled completely eliminated activity (Fig. 4B, C). The mutation H262A also completely abolished activity, suggesting a direct role in catalysis beyond its membrane interactions alone. This is discussed further below. Activity was nearly abolished in K208D and sharply reduced in T244A, with smaller but significant reductions seen in K62D/K64D and Y209A. Apparent reductions were seen in Y210A, R265A, and H266A, but did not rise to statistical significance. The observation that most of these mutations had at least some effect on catalysis in the SUV system confirms the predicted membrane interaction surface identified by the MD simulations.

**Figure 4.**
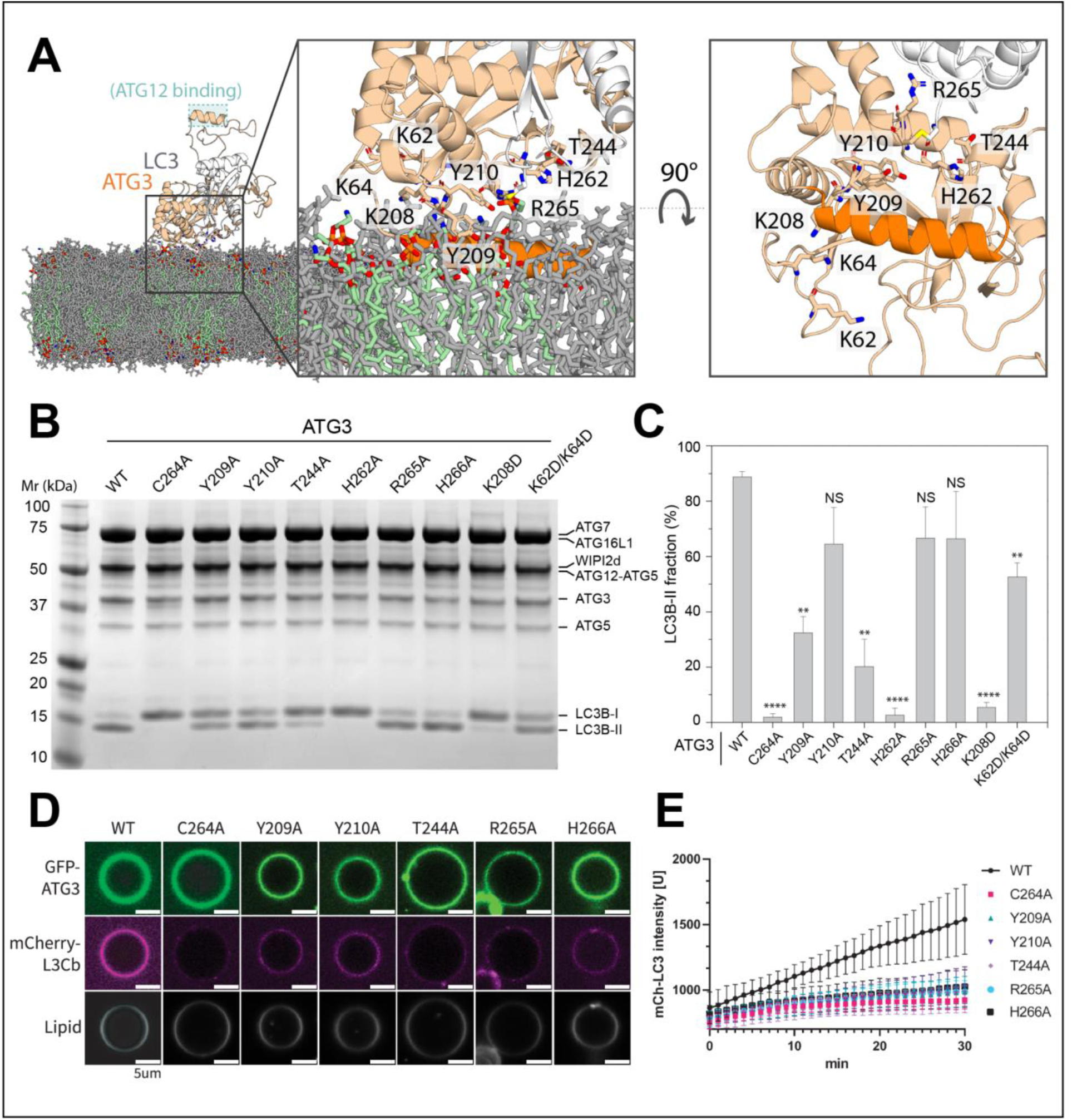
Mutational analysis of ATG3 membrane interaction face. (**A**) Snapshot of ATG3-LC3 upon spontaneous membrane association, taken at *t* = 1 μs of a 2 μs simulation replicate. Zoomed-in views from the side (left) and bottom (right; with lipids omitted for clarity) are shown. Phosphatidylethanolamine lipids are highlighted in green. (**B**) ATG3 *in vitro* LC3 lipidation. Do-SUVs [70% PC:20% PE:5% PS:5% PI(3)P] were incubated with ATG3 wild-type or mutant, ATG7, E3, WIPI2d, and LC3B. After 20 min, samples were loaded onto a 4-15% SDS-PAGE gel and stained with Coomassie blue. (**C**) Quantification of *in vitro* LC3 lipidation results, plotting the LC3B-II percentage in the total band intensities of LC3B-I and LC3B-II. P values were calculated using Student’s t-test: not significant (ns), P ≥ 0.05; **, 0.001 < P < 0.01; ***, P < 0.001. (**D**) Representative confocal images of GUVs showing GFP-ATG3 membrane binding and mcherry-LC3 lipidation. E3, WIPI2d, ATG7, GFP-ATG3 WT or mutant, mCherry-LC3B, and ATP/Mg^2+^ were incubated with GUVs (64.8% DOPC: 20% DOPE: 5% DOPS: 10% DOPI (3)P: 0.2% Atto647 DOPE) at room temperature. Images taken at 30 min are shown. Scale bars, 5 µm. (**E**) Quantification of relative intensities of mCherry-LC3B on GUV membranes (means ± SDs are shown; N = 30).

SUV assays are relatively permissive for LC3 lipidation by ATG3, likely due to the favorable contribution of ATG3 helix α1 to curved membrane binding^15^. Giant unilamellar vesicle (GUV) assays have a more stringent dependence on regulatory cofactors^17, 42^, so we repeated a subset of the above LC3B lipidation assays in this setting (Fig. 4D, E). Qualitatively, the results were similar, with C264A completely inactive, and others showing reduced activity. The rank order of the mutational effects subtly diverges from the SUV system, which is not surprising, given the differing physical presentation of the membrane to the conjugation complex. The main conclusion is that all of the mutations have at least some impact on activity in the GUV system, again consistent with the predictions of the simulations.

To determine if the predicted membrane function had the same function in living cells as in the reconstituted system, we generated an ATG3 KO HeLa cell line. ATG3 KO was verified by Western blotting (Fig. 5A). Starvation-induced autophagic flux was monitored with the HaloTag-LC3B system based on the appearance of a free HaloTag-band^17^. As expected, expression of the wild-type construct rescued autophagic flux in the KO cells, while no flux was observed in the C264A rescue (Fig. 5B, C). In a contrast to the partial effects seen for most mutants in vitro, an almost complete loss of flux was seen in most of the mutants in the ATG3 KO cells. This may reflect more stringency in the cellular system. H266A, which has a marginal (not statistically significant) reduction in activity in SUVs and a larger reduction in GUVs, has essentially no loss of activity in cells. The more variable effects of H266A in different assays may reflect a higher degree of context-dependence of the function of His266, as discussed below. The main conclusion from the ATG3 KO experiments is that the membrane interaction surface identified in the MD simulations accurately predicted loss of function in the cellular setting.

**Figure 5.**
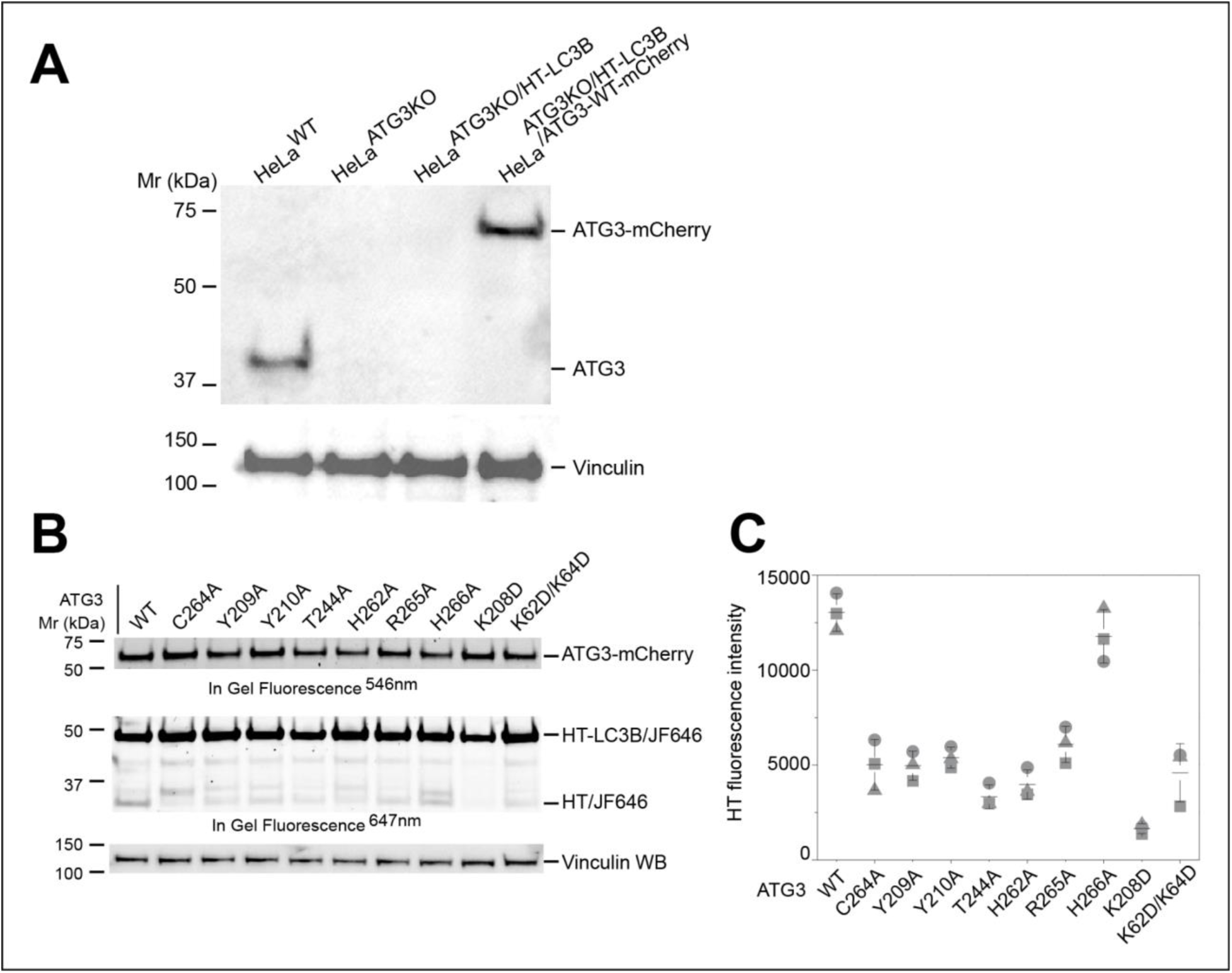
Mutations in ATG3 membrane interaction face impair function in cells. (**A**) ATG3 western blot of HeLa^WT^, HeLa^ATG3KO^ derivative cells. (**B**) HeLa^ATG3KO^ stably expressing HT-LC3B and ATG3-mCherry wild-type or mutant were starved in EBSS for 4 hours. Cells were pulse-labeled with 50 nM JF646-conjugated HT ligand before starvation. In gel fluorescence of mCherry and JF646 were imaged at 546 nm and 647 nm. (**C**) Quantification of free Halo^JF646^ band intensities of in-gel fluorescence. Mean values and standard deviations from three experiments are shown with a line and error bar.

### Conserved His262 of HPC motif facilitates ATG3-catalyzed LC3 lipidation

Previous studies have shown that the transfer of LC3 from ATG3 to lipid substrates is sensitive to pH and takes place more efficiently under slightly basic conditions *in vitro*, most likely through an effect on the ATG3 conjugase activity^47, 48^. Whilst the protonation state of the target PE amine group is expected to show little variation within the pH range of interest, we note the presence of two histidine residues, His262 and His266, in close proximity to the catalytic Cys264. Both histidines are fully conserved across ATG3 homologues and, with their characteristic pKa just below physiological pH, serve as possible acidity sensors for the ATG3-catalyzed reaction.

In atomistic MD simulations of the ATG3-LC3 conjugate with the His262 and His266 side chains both in their unprotonated state (which is predicted to be the dominant species at pH ≥ 7), His266 remained oriented towards the protein interior with a minimum distance of ∼1 nm to the nearest lipid (Fig. 6A). By contrast, frequent lipid interactions formed by His262 are suggestive of a direct role in the LC3 conjugation reaction. Whereas the nucleophile of the reaction, the PE amine group, did spontaneously approach the backbone carbonyl carbon of Gly120 (LC3) to be attacked (reaching a minimum distance of ∼0.4 nm), such interactions were infrequent. Meanwhile, the unprotonated nitrogen of the His262 imidazole was observed to interact with the positively charged primary amine of PE headgroups within bonding distance (< 0.2 nm) to the amine proton (Fig. 6A). Furthermore, our simulations capture a configuration in which the His262:PE interaction coincided with that between PE and Gly120 (Fig. 6B).

**Figure 6.**
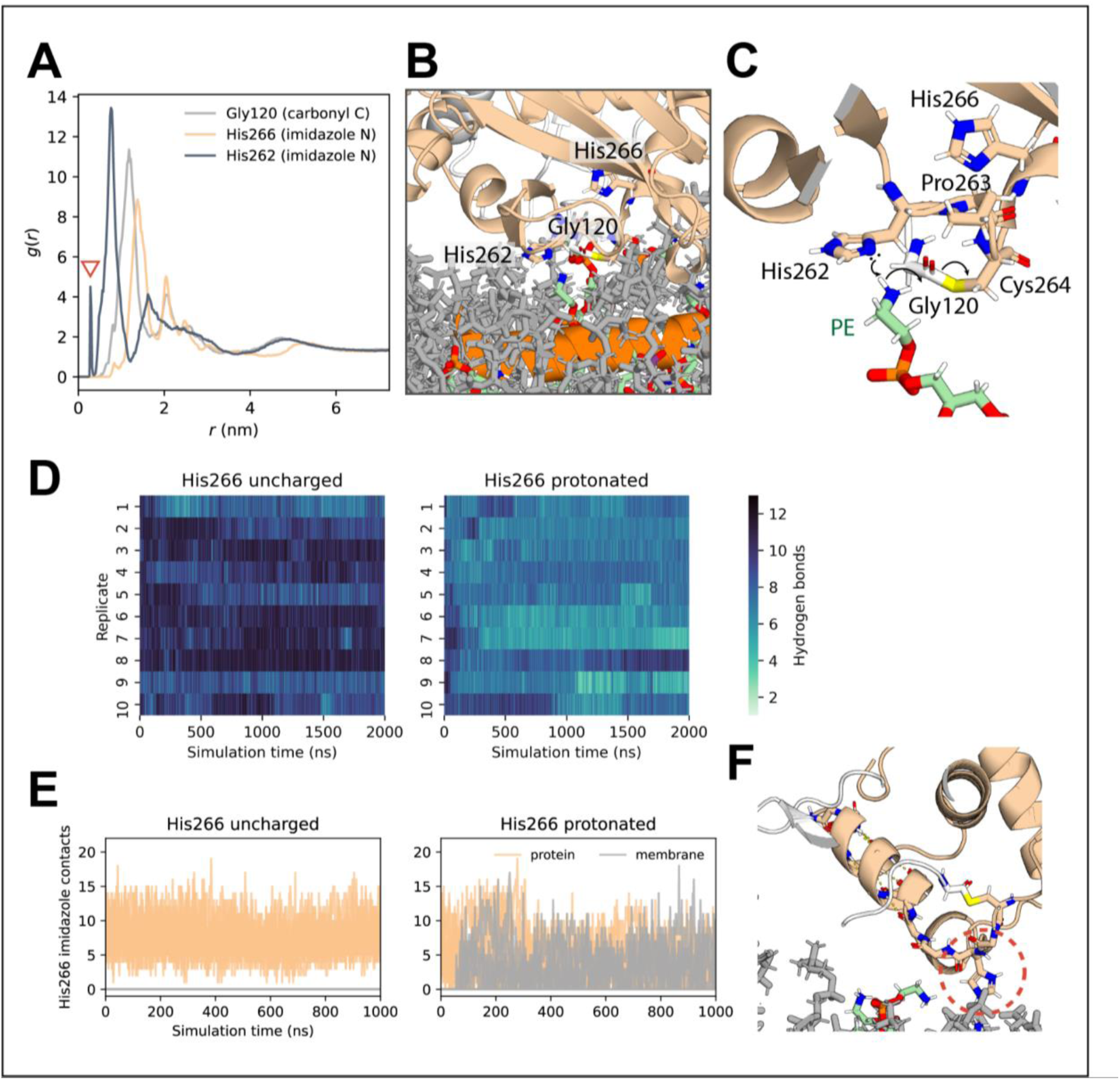
Two conserved histidine residues around ATG3 active site assume distinct roles. (**A**) Radial distribution function of protein atoms around the amine nitrogen atom of membrane phosphatidylethanolamine (PE) lipids (arrow: direct contact). (**B**) Snapshot of ATG3-LC3 interacting with membrane lipids during atomistic molecular dynamics simulation. The zoomed in view captures a PE lipid binding into the ATG3 active site, near the thioester bond under attack. The nearest PE amine proton also interacts within bonding distance (< 0.2 nm) of the unprotonated nitrogen atom of the His262 imidazole ring. (**C**) Possible mechanism for initiation of LC3 lipidation reaction, whereby the ATG3 His262 imidazole ring would facilitate a nucleophilic attack on the Gly120 carbonyl of LC3. The backbone amide of Cys264 is in position to stabilize the developing negative charge on the Gly120 oxygen. Illustrated using the simulation snapshot of (**B**), showing only the ‘attacking’ lipid for clarity. (**D**) Number of backbone hydrogen bonds in ATG3 helix 265-280 in simulations of ATG3-LC3 with ATG3 His266 in an uncharged or doubly protonated state. (**E**) Number of protein and membrane lipid contacts made by the His266 imidazole ring in the uncharged and the doubly protonated state, respectively, in simulations of alternative models of the ATG3-LC3 conjugate. (**F**) Snapshot of membrane-associated ATG3-LC3 in which the side chain imidazole of His266 (indicated with a red dashed circle) was doubly protonated with a charge of +1. At *t* = 1 μs of the 2 μs simulation replicate shown, destabilization of the local protein structure has brought the His266 side chain into membrane contact.

His262 and Cys264 of human ATG3 form part of the HPC motif that is conserved across orthologues of ATG3 as well as ATG10, an E2-like autophagic enzyme that catalyzes ATG12 conjugation to ATG5^6^. Combined with previous^35, 49^ and present evidence of a critical role of His262 for ATG3 conjugase activity, our simulation results are suggestive of a plausible reaction mechanism in which the His262 imidazole ring would deprotonate the PE amine group for nucleophilic attack on the Gly120 carbonyl of LC3 (Fig. 6C). As part of such a proposed mechanism, the unique backbone conformational restraints conferred by the cyclic side chain of Pro263 in the HPC motif would be crucial for orienting His262 and Cys264 side chains in relative positions conducive to catalysis, explaining their full conservation (SI Fig. 3). The protein backbone conformation conferred by Pro263 also holds the backbone amide of Cys264 within bonding distance of the carbonyl oxygen of Gly120 (Fig. 6C), which would stabilize the oxygen anion intermediate formed during the reaction. Energetically favorable breakage of the thioester bond will then yield the LC3-PE conjugate, a stable amide product. Alternatively, ATG3 has been reported to catalyze conjugation of ATG8 family proteins to phosphatidylserine (PS) lipids in the non-canonical pathway of autophagy^50^. In accordance with this, our simulations of ATG3-LC3 conjugate also capture an analogous membrane-interacting configuration likely poised for reaction with a PS molecule (SI Fig. 4).

His266, the second of the two conserved histidine residues described above, has been implicated in the pH-dependent conjugase activity of ATG3 in a recent study^35^. To assess the effect of altering the protonation state of His266, we performed additional MD simulations of the ATG3-LC3 conjugate, in which the His266 imidazole ring was doubly protonated. Strikingly, the extra proton destabilized the local protein structure (Fig. 6D). A reorientation brought the His266 side chain into direct membrane contact within the first hundreds of nanoseconds in seven out of ten simulation replicates (Fig. 6E, F). These results are consistent with His266 fulfilling a pH-sensitive structural role, as previously proposed for its counterpart in ATG3 orthologues (His236 in the yeast protein and His260 in *Arabidopsis*)^48^, and provide an explanation for the alternative conformations in this region between available crystal structures obtained at different pH^44, 51^.

## Discussion

Building upon an increasing collection of structural and biochemical data on the components and interactions that form the autophagic LC3 lipidation machinery, we set out to complete the molecular puzzle of how the E3-like ATG12–ATG5-ATG16L1 complex and the E2-like conjugase ATG3 deliver LC3 to phagophore membranes. Results from atomistic MD simulations point toward a multistage mechanism progressively localizing the ATG3-LC3 conjugate nearer to the target membrane and orienting the reactive center of LC3 conjugation toward lipid substrates. This process requires the sequential action of three previously identified membrane sensors within the assembly: (i) WIPI2 as the PI(3)P effector protein that drives membrane recruitment of ATG12–ATG5-ATG16L1^22, 52^, (ii) the curvature-sensitive ATG16L1 helix α2 within the (ATG12–)ATG5-binding domain^25, 26^, and (iii) the N-terminal amphipathic helix and membrane docking face of ATG3^43^.

As an emerging theme in cellular processes, with analogies to the multi-step process of docking in vesicle fusion^53^, the stepwise mechanism for the membrane targeting of LC3 provides additional layers of regulatory potential to the autophagic pathway. On the protein side, phosphorylation and other post-translational modifications will affect the stability, accessibility, and affinity of the distinct interaction elements. On the membrane side, variations in lipid composition and PI phosphorylation will modulate membrane recruitment. The phagophore lipid composition in particular modulates the recruitment of WIPI2 as anchor for ATG16L1 in docking step 1 as well as the membrane insertion of the ATG16L1 α2 helix and the ATG3 N-terminal helix in steps 2 and 3, respectively. Growing evidence points to a second WIPI2-interacting site within the ATG16L1 coiled-coil domain^27^, which would facilitate step 2 of our model in a PI(3)P-dependent manner. Occupancy of the second site for WIPI2 has been proposed to facilitate LC3 lipidation following the initial membrane recruitment of ATG12–ATG5-ATG16L1, in line with earlier observations of allosteric activation of the complex by WIPI2^42^. Consistent with our structural model, a second WIPI2 molecule bound to the coiled-coil region pulls the N-terminal side of ATG16L1 closer to the membrane surface (SI Fig. 5), thereby facilitating the membrane insertion of the ATG16L1 α2 helix in step 2 of our docking model.

Through multi-microsecond all-atom MD simulations, collecting a total of > 50 μs membrane docking trajectories, we have examined the lipid-interacting regions of the complete ATG3-LC3 conjugate for molecular determinants of the LC3 conjugation mechanism. The spontaneous membrane insertion of the ATG3 N-terminal helix in our simulations is consistent with a role of this region in positioning the protein onto the membrane for subsequent enzymatic activity^15, 54, 55^. Additional membrane-interacting residues concentrate around the Cys264 residue holding LC3. Mutations of these residues impact lipidation both *in vitro* and *in vivo*, confirming the catalytic relevance of the observed membrane interactions.

Simulations and experiments identify distinct roles for two fully conserved histidine residues in the vicinity of the catalytic cysteine of ATG3. We found neutral His266 to stabilize a catalytically competent structure of the active site, consistent with retained lipidation activity of the H266A mutant. By contrast, protonation of His266 disrupted the active site in our MD simulations, consistent with a role of His266 as pH sensor^35^. Whilst uncharged His266 serves to stabilize the catalytic loop conformation, our data point to active participation of His262 in the initiation of the LC3 lipidation reaction. In particular, we found the unprotonated His262 imidazole nitrogen to be positioned as proton acceptor from PE. Consistent with a possible catalytic role, the H262A mutation abolished function. His262 is the starting residue in the highly conserved HPC motif^45^ of ATG3, which is shared with the ATG10 conjugase family. However, in ATG10 the nearby His266 of ATG3 is changed to a threonine, which may reflect the distinct substrate specificity of the two enzymes (SI Fig. 3).

The critical biological role of the ATG12–ATG5-ATG16L1 complex in mammalian autophagy^19^, and before that, the role of the corresponding Atg12–Atg5-Atg16 complex in yeast^16^, has long been appreciated. Yet the precise role of this complex in LC3 lipidation has been hard to define. The role of the extensive structural elements linking the N-terminal helix of ATG3 on the one hand, and the established WIPI2-dependent membrane docking site on the other, have proven difficult to characterize as the membrane-associated system is too large for NMR, yet too dynamic for X-ray crystallography or single-particle cryo-EM. Under the “computational microscope” of molecular dynamics simulations, the role of the connecting elements in mediating a stepwise docking process has now been unveiled. As a core element in the molecular machinery of selective autophagy, this new and far more detailed insight into the membrane docking steps of LC3 will undoubtedly facilitate the therapeutic targeting of autophagy in Parkinson’s disease and other neurodegenerative diseases.

## Materials and Methods

### Structural models of protein complexes

Atomistic models of the ATG3-ATG12–ATG5-ATG16L1 and WIPI2d-ATG16L1 complexes were based on crystal structures with PDB IDs 4NAW^21^ and 7MU2^18^, respectively. The ATG16L1 N-terminal domain in the former complex was replaced by a more complete structure (PDB ID: 4TQ0^56^). As introduced previously^25^, an alternative conformation of the same ATG16L1 region was generated in PyMOL^57^ by rotation of helix α2 relative to helix α1 at the Gln30/Ala31 hinge. A model for the dimeric central ATG16L1 domain was completed through (i) homology modeling of residues 141-225 using SWISS-MODEL^58^ based on crystal structures of the mouse protein (PDB IDs: 6ZAY^32^ and 6SUR^31^) and (ii) parameter fitting for residues 78-193 with CCBuilder 2.0^29^ upon coiled-coil prediction^28^ by NPS@^59^. Unstructured inter-domain loops were added using the DEMO server^60^ to yield an ATG16L1 dimer encompassing residues 1-247. AlphaFold v2.2^34^ was used to model ATG3-LC3B in complex with the ATG7 homodimer. The Cys264 side chain of ATG3 was connected to the LC3B C-terminus by a thioester bond, parameterized using CHARMM-GUI^61,62^. The ATG5 Lys130 side chain was similarly connected to the ATG12 C-terminus, via an isopeptide bond. The ATG16L1 WD40 domain (dispensable for canonical autophagy^63^) was excluded from the model, as were the unstructured ATG12 residues 1-52 and WIPI2d residues 1-11 and 362-425. Exposed N- or C-terminal groups at the end(s) of each incomplete structure or truncated construct were neutralized. Protonation states of amino acid side chains were assigned according to pKa prediction by PROPKA^64^. His183 and His255 at the putative PI(3)P binding sites of WIPI2d were protonated. Six models of the ATG3-LC3B conjugate were generated, with the imidazole of ATG3 His262 uncharged (protonated at the δ- or ɛ-nitrogen in alternative models) and that of His266 uncharged (protonated at δ- or ɛ-nitrogen in alternative models) or cationic (doubly protonated).

### Molecular dynamics simulations

Molecular dynamics simulations were performed with GROMACS 2020^65^ using the CHARMM36m force field^66^. Following the same protocol as previously described^25^, all membranes consisted of 60% DOPC, 20% DOPE, 5% DOPS, 10% POPI^38^, and 5% PI(3)P based on the ER lipid composition^12^ and were prepared initially in a coarse-grained representation using the *insane* method^67^. Curved membranes were constructed using LipidWrapper^68^ by fitting the amplitude of the membrane buckle as a sine function of its *x*-coordinate. Each coarse-grained membrane system was solvated with 150 mM of aqueous NaCl, equilibrated for 200 ns, and converted into an atomistic representation using the CG2AT2^69^ tool. Atomistic models of protein complexes were placed above membranes after CG2AT2 conversion, followed by re-solvation and 10 ns of further equilibration. Simulation replicates were independently prepared and equilibrated. During equilibration, harmonic positional restraints with a force constant of 1000 kJ mol^-1^ were applied to non-hydrogen protein atoms or backbone beads. The *xy* dimensions of buckled membrane systems were fixed in simulations. System temperature and pressure were maintained at 310 K and 1 bar, respectively, using the velocity-rescaling thermostat^70^ and a semi-isotropic Parrinello-Rahman barostat^71^ during the production phase. The integration time step was 2 fs. Long-range electrostatic interactions were treated using the smooth particle mesh Ewald method^72,73^ with a real-space cut-off of 1 nm, a Fourier spacing of 0.12 nm, and charge interpolation through fourth-order B-splines. The LINCS algorithm was used to constrain covalent bonds involving hydrogen atoms^74^. Simulation trajectories were analyzed through MDAnalysis 2.0^75,76^ in Python 3.6.

### Plasmids

ATG3 wild-type and mutants were sub-clones into 1GFP or 2BT vector which contained an N-terminal 6His-GFP or 6His tag, respectively, using ligation independent cloning. Plasmids of pVSV-G (Addgene #138479), pBS-CMV-gagpol (Addgene #35614) and pMRX-IP-HaloTag7-LC3 (Addgene #184899) were used for retroviral generation of ATG3 KO/HaloTag7-LC3 stable cell line. Plasmids of VSV-G, pCMV R8.74 (Addgene #22036) and ATG3-mCherry viral vector were used for lentiviral generation of ATG3-mCherry wild type and mutants into ATG3KO/HaloTag7-LC3 cell line. The mCherry tag was inserted between ATG3 (1-125) and (126-314). PCR products of ATG3-mcherry wild-type and mutants were subcloned into pCDH1-CMV-MCS-SV40-hygromycin viral backbone using NheI and BamHI. All constructs were verified by DNA sequencing.

### Protein expression and purification

ATG3 constructs used for in-vitro lipidation assays, GUV assays were all expressed in *E. coli* (BL21) DE3 star cells. Cells were grown in Lura Bertani (LB) media at 37°C until an OD_600_ of 0.8. The culture was induced with 1 mM IPTG and grown overnight at 18°C. Cells were pelleted and resuspended in 50 mM HEPES pH 7.5, 300 mM NaCl, 2 mM MgCl_2_, 10 mM imidazole, 1 mM tris(2-carboxyethyl)phosphine [TCEP]) supplemented with EDTA-free protease inhibitors (Roche). The cells were lysed via sonication and lysate was clarified by centrifugation (17,000 rpm for 1 hour at 4°C). The supernatant was then applied to 1mL of Ni-NTA resin. The resin was subsequently washed thoroughly with at least 100CV of lysis buffer, and the protein was eluted with lysis buffer supplemented with 300 mM imidazole. The eluted proteins were concentrated and loaded onto a Superdex 200 column (10/300 GL; GE Healthcare) equilibrated with a buffer containing 25 mM HEPES, pH 7.5, 150 mM NaCl, and 1 mM TCEP. Peak fractions corresponding to the protein were collected, pooled, snap frozen in liquid nitrogen, and stored at –80°C. Purification of ATG12–ATG5-ATG16L1, ATG7, and LC3 used for liposome lipidation assays, GUV assays were performed as previously described^42^. Purification of WIPI2d was performed as previously described^18^.

### In-vitro LC3 lipidation assays

A lipid mixture with a molar composition of 70% DOPC, 20% DOPE, 5% DOPI(3)P, and 5% DOPS (Avanti Polar Lipids) was dried under a nitrogen stream and put under vacuum overnight. Lipids were resuspended at 1 mg/mL in the assay buffer (25 mM HEPES pH 7.5, 135 mM NaCl, 2.7 mM KCl, 1 mM TCEP), freeze thawed seven times, and extruded 17 times through a 100 nM filter (Whatman). Reactions were set up at room temperature in the assay buffer to a final concentration of 1 µM of the indicated ATG3 construct, 1 µM ATG7, 1 µM E3, 500 nM WIPI2d, 5 µM LC3B, 0.5 mM ATP, 1 mM MgCl_2_, and 0.5 mg/mL liposomes. 15 µl of Reactions were quenched at 20 minutes with 4xLDS loading buffer, boiled at 60°C for 10 min, and then loaded onto SDS PAGE gels. Protein bands were visualized with Coomassie blue. Three biological replicates were performed. Protein band intensity of LC3B-I and LC3B-II was analyzed by ImageJ (https://imagej.nih.gov/ij/). Quantification of LC3B-II formation was plotted as percentage of total LC3B among the measured values per each ATG3 protein in a bar graph. Averages and standard deviations were calculated. The P values were calculated using an unpaired two-tailed Student’s t test. P values were considered as follows: not significant (NS), P ≥ 0.05; *, 0.01 < P < 0.05; **, 0.001 < P < 0.01; ***, 0.0001 < P < 0.001; and ****, P < 0.0001.

### GUV assay

GUVs were prepared by hydrogel-assisted swelling as described previously^17^. Briefly, A lipid mixture with a molar composition of 64.7% DOPC, 20% DOPE, 10% DOPI(3)P, 5% DOPS and 0.3% Atto647N DOPE was dried onto a PVA film and put under vacuum overnight. GUVs were swelled with 300mM sucrose at room temperature for 1 hour. The reactions were set up in an eight well observation chamber (Lab Tek) that pre-coated with 1 mg/ml bovine serum albumin (BSA). The reactions were mixed at room temperature to a volume of 120 uL in 25 mM HEPES pH 7.4, 150 mM NaCl, 1 mM TCEP. A final concentration of 100 nM ATG3-GFP or mutant proteins, 200 nM WIPI2d, 50 nM E3 complex, 100 nM ATG7, 100 nM ATG3, 500 nM mCherry-LC3B, 5 mM ATP, and 1 mM MnCl_2_ were used. 3 µL GUVs were added last to initiate the reaction. After 5 min incubation, during which random views were picked for imaging, time-lapse images were acquired in multitracking mode on a Nikon A1 confocal microscope with a 63 × Plan Apochromat 1.4 NA objective. Three biological replicates were performed for each experimental condition. Identical laser power and gain settings were used during the course of all conditions.

For quantification of protein intensity on GUV membranes, the outline of individual vesicle was manually defined based on the membrane channel. The intensity threshold was calculated by the average intensities of pixels inside and outside of the bead and then intensity measurements of individual GUV were obtained. Averages and standard deviations were calculated among the measured values per each condition and plotted in a bar graph. The data were analyzed with GraphPad Prism 9.

### Mammalian cell culture

HEK293T and Hela cells were cultured in Dulbecco’s Modification of Eagle’s Medium (DMEM) with GlutaMAX (Gibco 10566016) supplemented with 10% fetal bovine serum (FBS) and 1% Penicillin-streptomycin in a 5% CO_2_ incubator at 37°C.

### Retroviral/lentiviral generation of stable cell lines

HEK293T cells were transfected with retroviral packaging plasmids (pVSV-G, pBS-CMV-gagpol) and pMRX-IP-HaloTag7-LC3 using TransIT-LT1 (Mirus). The viral supernatants were collected 72 hr after transfection followed by virus concentration using Lenti-X concentrator (TaKaRa). Virus containing media was added to ATG3 KO HeLa cells with 8 μg/mL polybrene (Sigma-Aldrich, H9268). After a day, selection was performed with 2 μg/mL puromycin (GoldBio, P-600). The protocol for generating ATG3 KO Hela cells stably expressing HaloTag-LC3B is available at <u>dx.doi.org/10.17504/protocols.io.5qpvo3xq9v4o/v1</u>.

To prepare the lentiviral solution, HEK293T cells were transfected with pVSV-G, pCMV R8.74 and pCDH1-CMV-ATG3-mCherry-SV40-Hygromycin using TransIT-LT1 reagent (Mirus). After 72 hr post-transfection, the lentiviral-containing media were collected, concentrated and then added into ATG3 KO/HaloTag7-LC3 HeLa cells with 8 µg/ml polybrene. After one day, selection was performed with 0.5 mg/ml hygromycin B (Thermo Fisher, 10687010). Positive clones were later selected by FACS sorting based on mCherry intensity using Wolf G2 cell sorter (Nanocellet). The protocol for generating stable ATG3-mCherry Hela cells with HaloTag-LC33B is available at <u>dx.doi.org/10.17504/protocols.io.dm6gp3exjvzp/v1</u>.

### ATG3 KO cell line generation

ATG3 KO cells (SI Table 1) were generated using a CRISPR guide RNA (gRNA) targeting Exon 5 of the ATG3 gene. Oligonucleotides (Sigma) that contain CRISPR sequences were annealed and ligated into BbsI-linearized pSpCas9(BB)-2A-GFP vector^77^ (a gift from Feng Zhang; Addgene plasmid # 48138). Sequence-verified gRNA constructs were then transfected into HeLa cells for 24 h and GFP-positive cells were individually sorted by fluorescence activated cell sorting (FACS) into 96 well plates. Single cell colonies were screened for the loss of the targeted gene product by immunoblotting. The PCR products generated from sequencing primers that anneal to the flanking regions of the targeting CRISPR (SI Table 2) in both the ATG3 KO and the WT parental line were directly sequenced. The resulting sequencing data was analysed using Synthego ICE v2 CRISPR Analysis Tool (https://ice.synthego.com/<u>#/</u>) (SI Table 1).

### HaloTag-LC3B processing assay

HaoTag-based assay was performed as previously described^78^. Cells were seeded at 100K Cells/well in 12-well plate one day before. Next day, cells were incubated in complete DMEM medium with 50 nM JF549 (Promega, GA1120) for 1 hr, and then washed twice with 1x PBS buffer followed by incubation with EBSS buffer (Thermo fisher, 24010043) to induce autophagy by starvation. After 4 hr starvation, cells were harvested with Trypsin (Thermo fisher, 25300120).

Cell pellets were washed with PBS and resuspended in lysis buffer (20 mM HEPES pH 7.5, 200 mM NaCl, 2 mM MgCl_2_, 10% glycerol, 1 mM TCEP, 0.2% *n*-dodecyl-β-D-maltoside (GoldBio, DDM25) with protease inhibitors (Thermo Fisher, A32963). 2 µl Benzonase (Merck Millipore, 70664) was added into each 500 µl lysis buffer. After 30 min of incubation on ice, cell lysates were centrifuged at 21,000g x 10 min, and then measured protein concentration using nanodrop spectrophotometer (Thermo Fisher).

For each sample, 20 µg clarified lysates were loaded onto NuPAGE 4-12% Bis-Tris Gel (Thermo Fisher, NP0322BOX). For in-gel fluorescence imaging, the gel was immediately visualized with ChemiDoc MP imaging system (Bio-Rad) after SDS PAGE. Band intensities were acquired by exciting samples at 546 nm and 647 nm. The protocol of HaloTag-LC3B processing assay to assess autophagy is available at <u>dx.doi.org/10.17504/protocols.io.3byl4qexzvo5/v1</u>.

## Acknowledgments

We thank members of the Aligning Science Across Parkinson’s Team mito911 for advice and discussions.

## Funding

The Michael J. Fox Foundation for Parkinson’s Research (MJFF) and Aligning Science Across Parkinson’s (ASAP) initiative. MJFF administers the grant ASAP-000350 on behalf of ASAP and itself. (J.H.H., M.L., G.H.)

Max Planck Society (S.R. and G.H)

## Author contributions

Conceptualization: JHH, GH Investigation: SR, LM, XR Formal Analysis: SR, LM, XR Visualization: SR

Resources: MS Supervision: ML, JHH, GH

Writing—original draft: SR, JHH, GH Writing—review & editing: SR, LM, XR, JHH, GH

## Data availability

Full data from molecular dynamics simulations (<u>10.5281/zenodo.8083724</u>) and from GUV experiments (<u>10.5281/zenodo.7988132</u>) have been uploaded at Zenodo. All data are available in the main text or the supplementary materials.

