## Supplementary Information for "Three-step docking by WIPI2, ATG16L1 and ATG3 delivers LC3 to the phagophore"

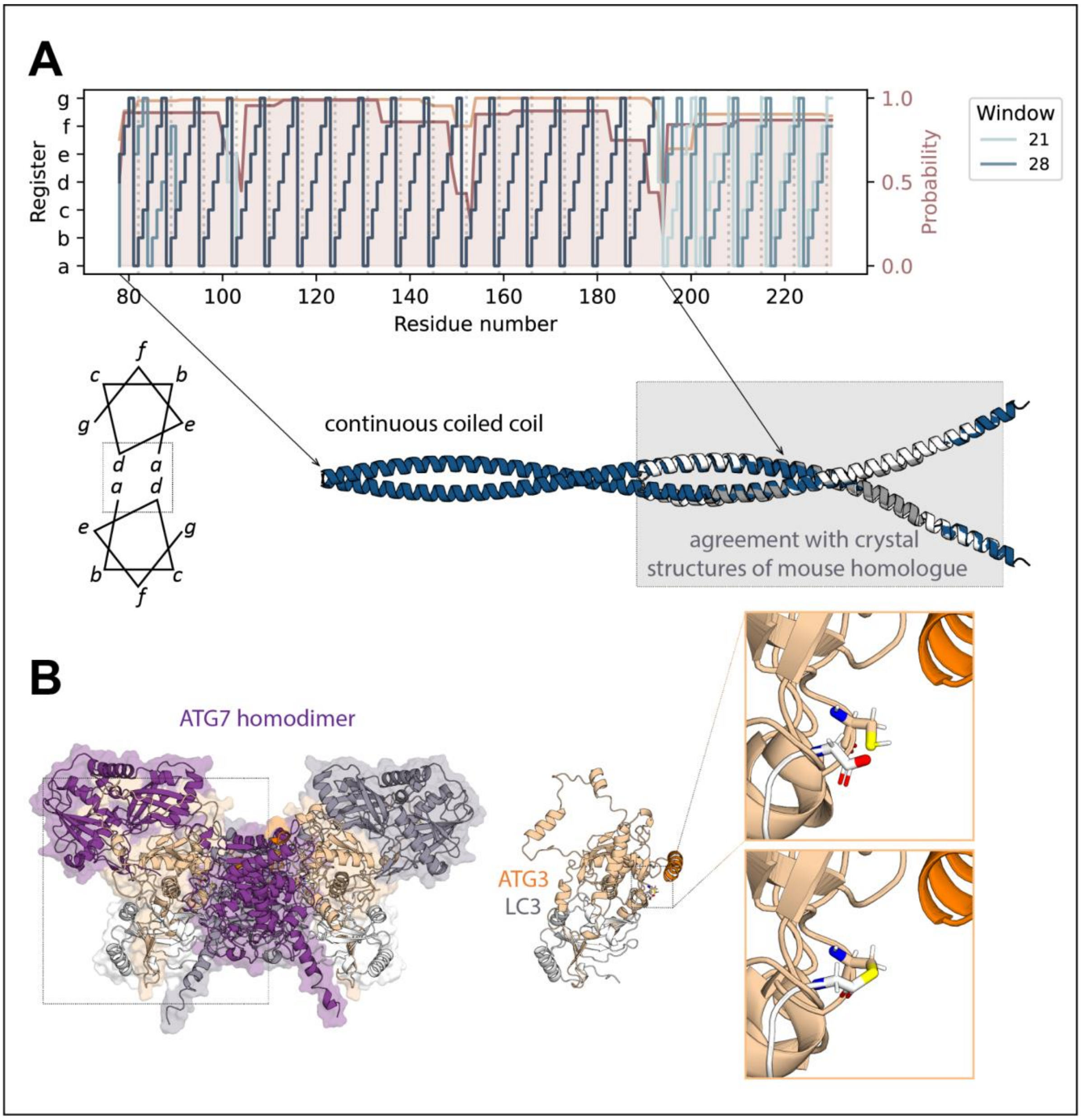


**Figure S1. Structural modeling of the human LC3 lipidation machinery components.** (**A**) Coiled-coil prediction and modeling of the central homodimerization domain of ATG16L1. The C-terminal region of the predicted stretch of continuous coiled coils (residues 78-193) is in excellent agreement with homology models based on crystal structures of the mouse orthologue (PDB IDs: 6ZAY and 6SUR, shown in white and gray, respectively) in which an overlapping region has been resolved. (**B**) Structure of ATG7 homodimer in complex with two copies of ATG3-LC3, as predicted by AlphaFold. As part of the larger complex, the catalytic Cys264 of ATG3 is in close proximity to the C-terminal Gly120 of LC3. A rotameric modification of the Gly120 conformation allows a thioester bond to be modeled, yielding the ATG3-LC3 conjugate.

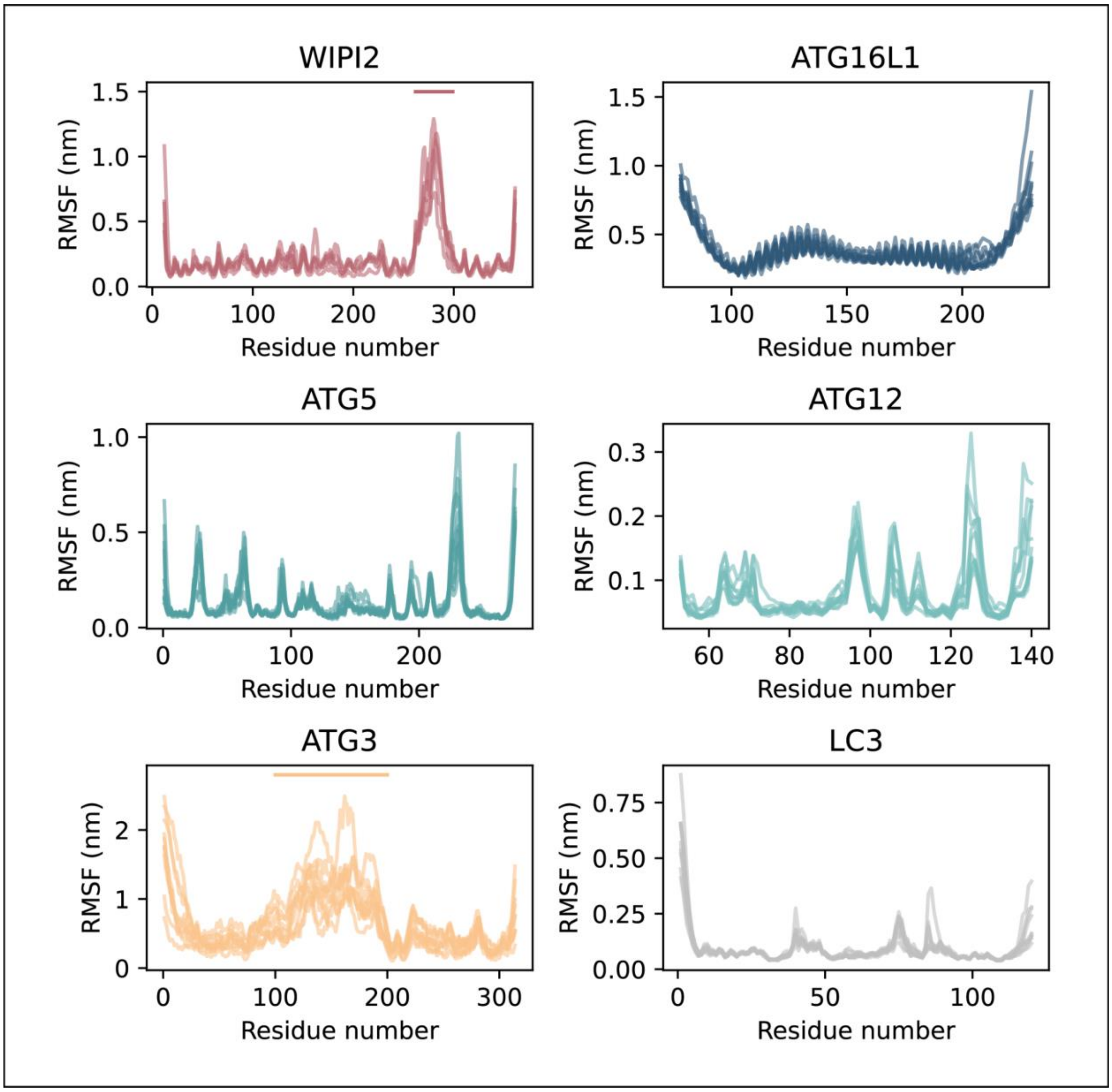


**Figure S2. Root-mean-square fluctuation (RMSF) assessment of protein stability.** Cα RMSFs of the membrane-associated ATG12–ATG5-ATG16L1-WIPI2 complex loaded with ATG3-LC3 are plotted for the respective constituent subunits during five 1 µs simulation replicates. Extended loop regions along the WIPI2 and ATG3 sequence are each indicated with a horizontal line.

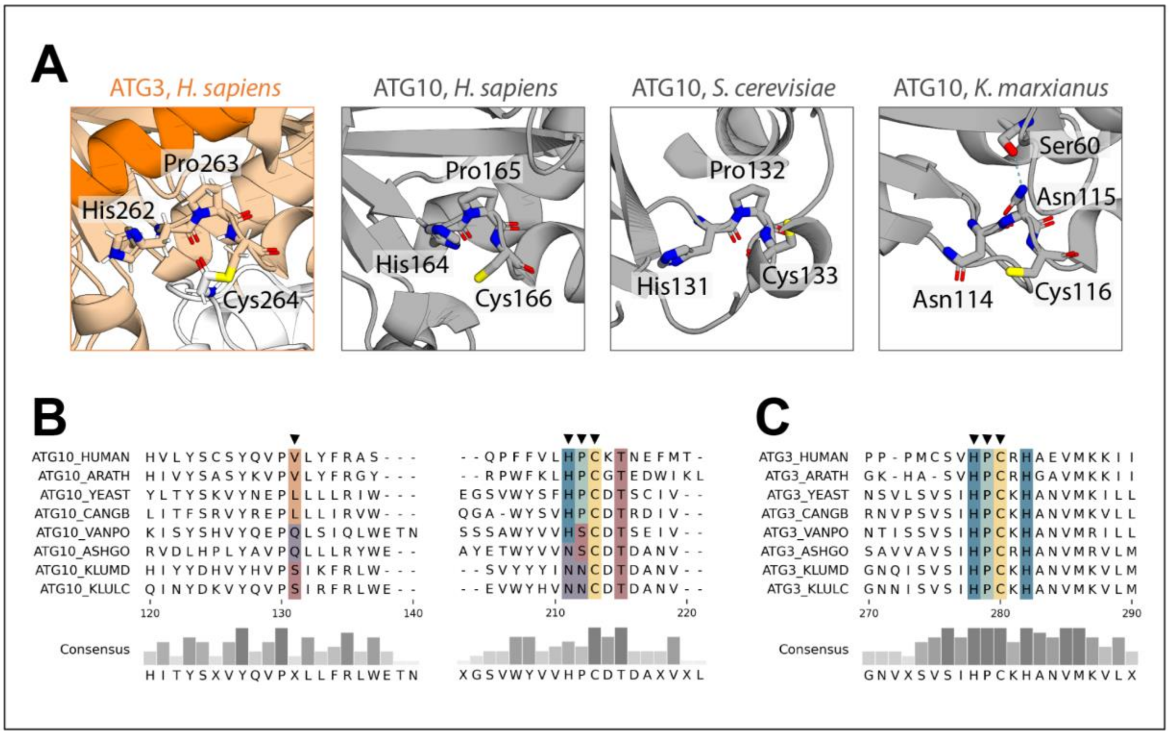


**Figure S3. Conservation of the HPC motif in the active site of ATG3 and ATG10 conjugase enzymes.** (**A**) Structure of the human ATG3 His262, Pro263, and Cys264 motif and its counterparts in the predicted structure of human ATG10 (from AlphaFold) and in crystal structures of two fungal homologs (PDB IDs: 4EBR and 3VX7), respectively. In each case, the cysteine and histidine (or asparagine) of the motif are held in the same relative orientation by the peptide backbone, stabilized either by the unique cyclic side chain of the proline or, in the case of the *K. marxianus* protein, through additional hydrogen bonding. (**B**) Sequence alignment of ATG10 orthologs reveals that substitution of the HPC motif proline by a serine or the larger asparagine residue is invariably accompanied by a corresponding mutation (from a hydrophobic residue to a glutamine or the smaller serine, respectively) at a position further upstream. The resulting formation of a serine-glutamine or asparagine-serine hydrogen bond would stabilize the peptide backbone of the motif in place of a proline. (**C**) Sequence alignment of ATG3 orthologs from the same selection of organisms.

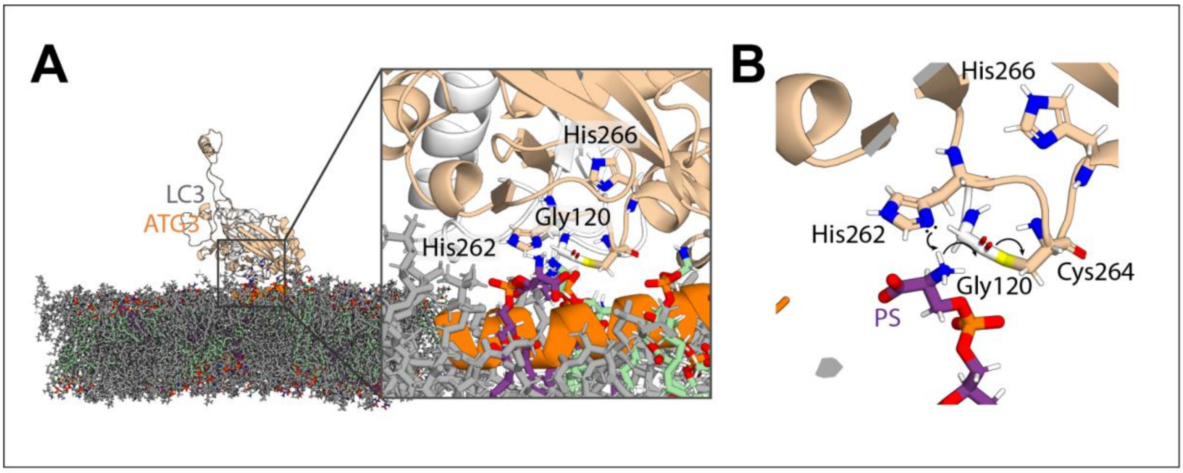


**Figure S4. Configuration of ATG3-LC3 conjugate poised for reaction with phosphatidylserine (PS) in atomistic molecular dynamics simulation.** (**A**) Snapshot of ATG3-LC3 interacting with membrane lipids, capturing a PS lipid binding into the ATG3 active site, near the thioester bond under attack. The nearest PS amine proton also interacts within bonding distance (< 0.2 nm) of the unprotonated nitrogen atom of the His262 imidazole ring. (**B**) Possible mechanism for initiation of LC3 lipidation reaction, whereby the ATG3 His262 imidazole ring would facilitate nucleophilic attack on the Gly120 carbonyl of LC3. The backbone amide of Cys264 is in position to stabilize the developing negative charge on the Gly120 oxygen. Illustrated using the same simulation snapshot from (**A**), omitting all lipids but one for clarity.

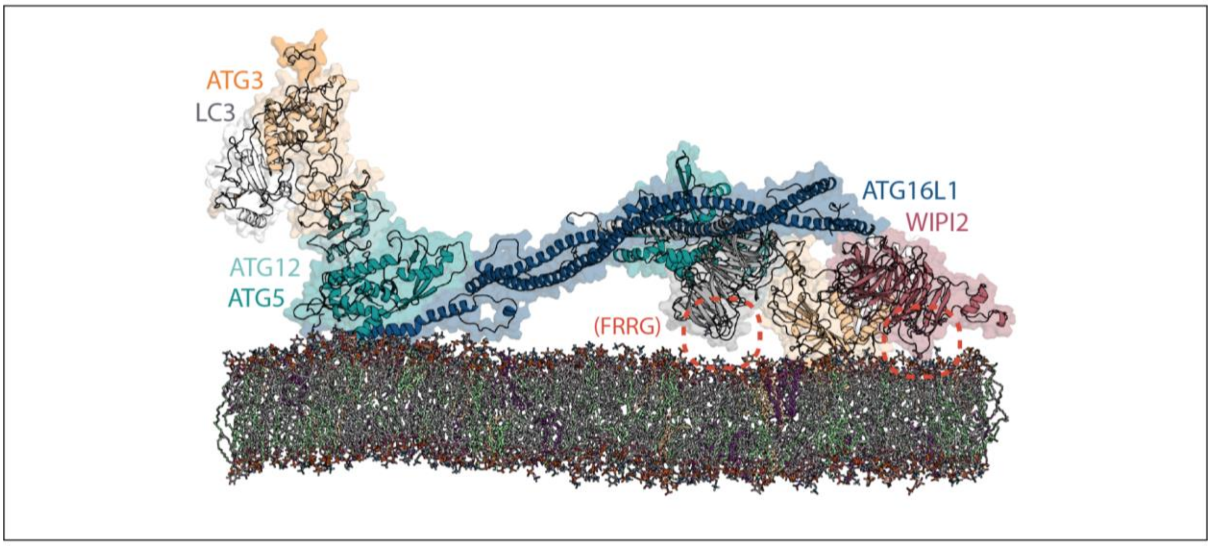


**Figure S5. Binding of a second WIPI2 to interaction site within the ATG16L1 coiled coil is compatible with membrane-engaged configuration of ATG12–ATG5-ATG16L1 complex from atomistic molecular dynamics simulation.** Locations of the membrane-interacting FRRG motif of the first (pink) and second (gray) WIPI2 molecule are each indicated with a red dashed circle.

**SI Table 1. ATG3 CRISPR sequence and genotyping results of ATG3 KO line**

| Gene Symbol | Uniprot | GeneID/Location (NCBI) | Exon targeting location | CRISPR gRNA (PAM) | Clone number used | Detected Alleles | Mutation | Protein Impact |
|---|---|---|---|---|---|---|---|---|
| ATG3 | Q9NT62 | 64422/NC_000003.12 | Exon 5 | TGTTTGCACCGCTTATAGCACGG | #24 | 2 | c.(243_244)insN[2];(243_244)insN[1] | p.(Y82?fs*28);(Y82?fs*2) |

**SI Table 2. Primers used for genotyping of ATG3 knockout lines.**

| Gene | Exon | Primer | Primer sequence (5'-3') | PCR product size (bp) |
|---|---|---|---|---|
| ATG3 | 5 | Forward | GAGCTGAATGTAACTCTTAATCACC | 378 |
| | | Reverse | TGGGACAAAACTATGCCTTATGT | |
